# Effects of visual feedback on corticostriatal activity during continuous *de novo* motor skill learning

**DOI:** 10.1101/2023.08.29.555448

**Authors:** Sungbeen Park, Junghyun Kim, Sungshin Kim

**Affiliations:** Department of Artificial Intelligence, Hanyang University, 04763 Seoul, Republic of Korea; Department of Data Science, Hanyang University, 04763 Seoul, Republic of Korea; Center for Neuroscience Imaging Research, Institute for Basic Science, Suwon 16419, Republic of Korea

**Keywords:** Striatum, motor skill learning, fMRI, visual feedback

## Abstract

A corticostriatal network plays a pivotal role in visuomotor learning. However, it is unclear to what extent visual feedback for motor learning impacts corticostriatal activity. To address this, we conducted an fMRI experiment in which participants acquired a complex motor skill using either online or offline visual cursor feedback. We found a highly selective response in the entire striatum to visual performance feedback in both conditions and a relatively higher response in the caudate nucleus for the offline feedback. However, the ventromedial prefrontal cortex (vmPFC) response was significant only for online feedback. Furthermore, we also found functional distinctions of the striatal subregions in random versus goal-directed motor control. These findings collectively underscore the substantial impact of such simple visual feedback, as designed in our study, in eliciting corticostriatal responses, thereby elucidating its significance in reinforcement-based motor learning.

## Introduction

Acquisition of a new motor skill entails an adaptive process of learning a mapping between motor commands and desired sensory outcomes, which serves as sensory feedback, including visual and proprioceptive feedback. Among the two types of feedback, visual feedback is a critical component of visuomotor learning, which guides motor planning to achieve a goal of a motor task, encouraging faster and more accurate performance ^1^. In motor adaptation tasks, for instance, the visual feedback can provide altered sensory outcomes from a perturbed environment, which are used to quantify performance error. Several studies have identified a cortico-cerebellar network as the neural substrate of the performance error, a learning signal in motor adaptation ^2-4^.

Visual feedback can also provide an evaluation of motor performance. In this context, the significance of this role of visual feedback becomes evident in the acquisition of novel motor skills, distinguishing it from the process of readjusting proficient movements, as seen in motor adaptation ^1,5,6^. In an alternate perspective, visual feedback functions as a conduit for transmitting the value associated with executed movements and sensory prediction errors, thereby facilitating the driving force behind reinforcement learning. In contrast to the cortico-cerebellar network for motor adaptation, a cortico-striatal network has been identified as a neural substrate for learning a new motor skill ^5,7,8^. Thus, visual feedback serves as a crucial learning signal by presenting sensory outcomes of motor commands, which affect distinct neural substrates involved in the learning process. However, most studies have focused on neural plastic changes following motor learning while ignoring the specific roles of visual feedback in motor learning ^9^. Consequently, it has been unclear how visual feedback contributes to learning and links to neural responses. Here, we present an fMRI experiment to examine how visual feedback influences learning a new motor skill. We sought to understand the effects of two distinct types of visual feedback—continuous (online) and discrete (offline) feedback involving the presence of a cursor on the screen or performance evaluation—on corticostriatal activity. Earlier studies similarly manipulated visual feedback to distinguish between neural substrates involved in internally and externally driven movements^10-12^. However, these studies employed a simple motor control task without learning a visuomotor mapping, thereby neglecting to examine how different types of visual feedback related to learning could elicit the corticostriatal activity.

For this, we employed a GLM analysis with a parametric regressor encoding time-varying learning performance. Then, we assessed the selectivity of the striatal activity to the visual feedback and further analyzed distinct activity patterns in the striatal subregions. We also contrasted the corticostriatal activity under online versus offline visual feedback condition. Finally, we analyzed the dataset from a separate experiment to differentiate between striatal activity related to goal-directed motor control and those linked to random motor control. In this experiment, participants executed a simple motor control task without any visual feedback, eliminating the learning aspect.

## Materials and Methods

### Participants

The study included twenty-six healthy young volunteers who were right-handed according to the Edinburgh Handedness Inventory ^13^ and had no history of neurological or psychiatric issues. Twenty-four participants (ten males, fourteen females; mean age = 24.9 ± 4.7 years, age range = 18-35 years) completed all fMRI task sessions. Two participants who claimed severe fatigue during the experiment were excluded from further analysis. All participants had a normal or corrected-to-normal vision and provided written consent in accordance with the dictates of the trust ethics committee of Sungkyunkwan University, Suwon, Republic of Korea. Participants underwent two scanning sessions for 1.5 hours at a 3T fMRI scanner as follows.

### Localizer session

A separate experiment was conducted to identify the region related to finger movement. When the visual cue “Move” was displayed, participants, equipped with an MR-compatible data glove (5DT Data Glove 14 Ultra), were directed to perform natural-speed movements with their right fingers, ceasing upon presentation of the “Stop” cue. Each “Move” or “Stop” phase persisted for 48 seconds, separated by 2-second intervals, and a total of six sets of “Move” and “Stop” conditions were executed, amounting to approximately 600 seconds in total. Then we recalibrated a mapping matrix **A** and an offset r_0_ using the finger-movement data from the last two “Move” blocks. Additionally, we ensured that all 25 grid cells on the 5 × 5 grid were reachable by finger movements (Figure 1A).

**Figure 1.**
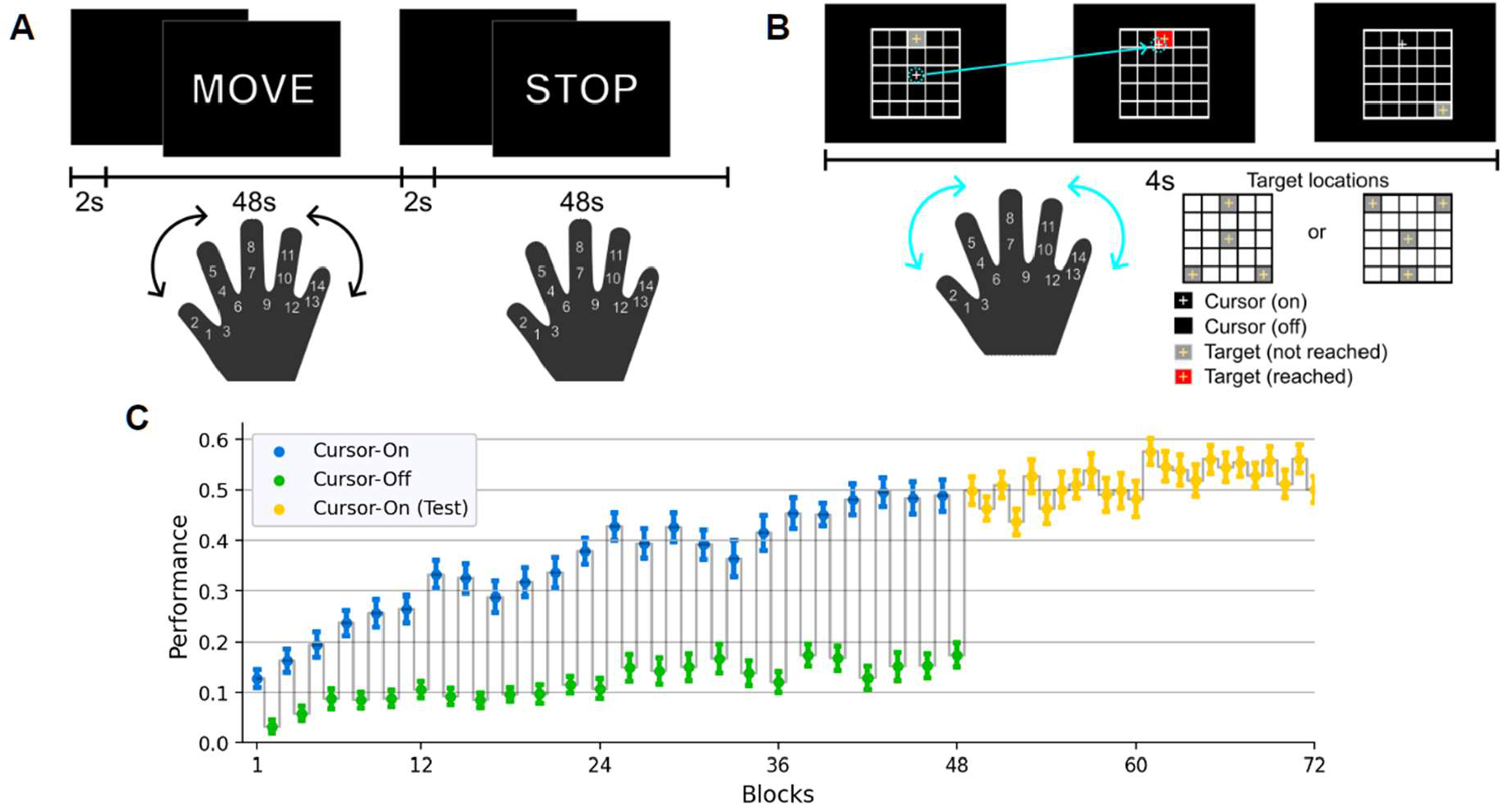
Experiment design and behavioral performance of all participants. (A) In the localizer session, participants move or stop their right fingers freely in response to the message displayed on the screen. The first principal components were calculated from the random finger movement by PCA and used as a finger-to-cursor movement mapping in the main session. (B) In the main session, a target appeared as a gray cell in the same order for each block, and participants were instructed to reach a target by moving their fingers. A target turned red when it was reached by a cursor which was visible in odd-numbered blocks (cursor-on condition) but not in the even-numbered blocks. Target locations were altered in the test condition where a cursor was always visible. (C) Block-by-block group performance. Participants improved performance across alternating “cursor-on” (blue) and “cursor-off” (green) learning blocks and continued in the test blocks (yellow). Error bars indicate SEM.

### Main session

We designed a task-based fMRI experiment consisting of a total of six runs using the same equipment as our previous study^14^. The mapping between hand postures and cursor positions was defined below.

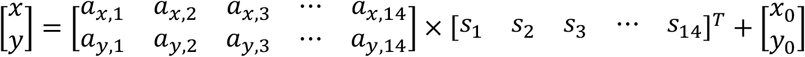

where *s*_*k*_(*k* = 1,2, …,14) indicates each of the 14 sensor inputs from the data glove, and *x* and *y* indicate the horizontal and vertical position of the cursor. In essence, the participants are required to acquire the proficiency in executing hand gestures, enabling the displacement of the cursor to the desired coordinates (x, y) throughout the task’s execution. The above equation can be rewritten as **r** = **As + r**_**0**_, where the mapping matrix A and the offset **r**_**0**_ were determined from the localizer session. Specifically, the mapping matrix’s first and second rows were determined as the first two principal components of the covariance matrix calculated from the last three blocks of movement data (“Move” condition) of the localizer session where participants made random finger movements.

The first part of the session consisted of four runs, each consisting of 144 trials from the current location to the target (twelve blocks of 12 trials). For 4 s during each trial, a gray grid cell with a yellow crosshair in its center appeared in one of the four target cells of a 5 × 5 grid, which is shown to participants (Figure 1B). Overall, the duration of each run is about 576 s (144 trials per run × 4 s per trial) without breaks between trials. For counterbalance, target cells were assigned in a triangular configuration for half of the participants or inverted triangular for the other half for the first main session. Given that the cell number is defined as *k* = 5*i* + *j* − 5, where *i* is a row index, *j* is a column index, and *i, j* ∈ ℕ), the target sequence in each block was ordered as cells **13-3-25-21-13-25-3-21-25-13-21-3** (triangle: sequence 1) or **13-23-5-1-13-5-23-1-5-13-1-23** (inverted triangle: sequence 2). This sequence was repeated for all twelve blocks during each run (144 trials) (Figure 1B).

The cursor position was continuously represented by a white crosshair in each run’s odd-numbered blocks (blue: On), while it was hidden in each run’s even-numbered blocks (green: Off) (Figure 1C). Meanwhile, when the cursor reached the target cell regardless of its visibility, the target cell changed to red (Figure 1B). Once the target was hit, participants were instructed to maintain a static position to remain in roughly the same location. The proportion of time during which the target turned red was measured as a trial-by-trial success rate. Thus, the goal of the task was to place the cursor on the target grid as quickly and precisely as possible and keep it there. Moreover, participants were instructed to move the cursor between targets as straight as possible. In the second part of a main session consisting of two runs, a target sequence was altered to the other sequence (sequence 1 or 2) (Figure 1B). The cursor position was consistently provided during these two runs (Test condition in Figure 1C).

#### fMRI Data Acquisition

We collected fMRI data using a 3-T Siemens Magnetom Prisma scanner with a 64-channel head coil. Functional images were acquired utilizing an echo planar imaging (EPI) sequence with the following parameters: 300 volumes (310 volumes for localizer fMRI); repetition time (TR) = 2,000 ms; echo time (TE) = 35.0 ms; flip angle (FA) = 90°; field of view (FOV) = 200 mm; matrix, 101 X 113 X 91 voxels; 72 axial slices; slice thickness = 2.0 mm. For anatomical reference, a T1-weighted anatomical scan of the entire brain was performed using a magnetization-prepared rapid acquisition with gradient echo MPRAGE sequence with the following parameters: TR = 2,300 ms; TE = 2.28 ms; FA = 8°; FOV = 256 mm; matrix = 204 X 262 X 260 voxels; 192 axial slices; and slice thickness = 1.00 mm. Prior to the functional scans, two EPI images were acquired with opposite-phase encoding directions (posterior-to-anterior and anterior-to-posterior) for subsequent distortion correction.

### Behavioral Data Analysis

MATLAB (version R2022a, MathWorks), Python (version 3.8.8), and Jupyter notebook (version 3.9.6) were utilized for all statistical analysis and data visualization. We used two-sided paired t-test between different experimental conditions. A trial-by-trial success rate was computed as a proportion of the time the cursor was on the target, i.e., targets turned on in red. As each task block included all 12 possible paths between four target locations and the same target sequence was repeated, we calculated an averaged success rate in each block, as shown in Figure 1C.

### fMRI Data Analysis

AFNI (Analysis of Functional NeuroImages, NIH; https://afni.nimh.nih.gov), MATLAB (version R2022a, MathWorks), Python (version 3.8.8), and Jupyter notebook (version 3.9.6) were utilized for all statistical analysis and data visualization.

### Preprocessing

Utilizing AFNI, anatomical and functional imaging data were preprocessed. Initially, localizer and task-based functional images were adjusted for slice-time acquisition and motion-related artifacts. The images were spatially registered to the anatomical data and translated into the Montreal Neurological Institute (MNI) template. All images were spatially smoothed using a Gaussian kernel with a 4 × 4 × 4-mm full-width at half-maximum, and the time series were scaled to have a mean of 100 and a range between 0 and 200.

### Whole-brain voxel-wise GLM analysis

To recognize regions modulating participants’ performance, “success” (i.e., reaching a target), we applied a parametric regressor (AFNI’s *3Ddeconvolve* with “stim_times_AM2” option) for a subject-level general linear model (GLM) analysis using six fMRI runs from the main task session. We first calculated the time-varying success rate for each trial of 4s-long time bins, defined as the proportion of time the cursor remained on the target grid cell and turned red. Then, we generated a pulse regressor whose amplitude is equal to the success rate at the offset of each trial and convolved it with a two-gamma function modeling a canonical hemodynamic response function (HRF). Then, we multiplied it with binary indicators coding for each feedback condition and generated two regressors of interest in the GLM analysis. However, for the result shown in Figure 3, we used a single regressor encoding all the trials. To localize regions related to finger movement, we first applied boxcar regressors encoding 48-s long “Move” and “Stop” conditions for a subject-level GLM analysis. Then, we calculated a whole-brain contrast map between the beta maps for the conditions, *β*_*move*_ > *β*_*stop*_.

For each of the fMRI runs, in both GLM analyses, five regressors modeling up to quartic (i.e., fourth-order) polynomial trends of the fMRI data and six regressors estimating rigid-body head motion were incorporated as nuisance regressors. The analysis excluded the volumes related to excessive head motion (defined as a displacement larger than 0.4 mm). After the voxel-wise whole-brain GLM analyses were performed for each subject, the group-level t-test was performed using AFNI’s *3dttest++* with the “-Clustsim” option, which is a conservative non-parametric Monte Carlo simulation method to determine cluster-corrected significance level. The voxel-wise threshold was set to p *<* 0.001, and the simulation determined 180 suprathreshold voxels as a cluster-wise corrected threshold p *<* 0.01 within the whole-brain group mask used in Table 1. The mask was defined as the intersection of all participants’ whole-brain scans by using *3dMean* with “mask_inter” option.

**Table 1.** Significant clusters positively modulating performance.

|  | Peak (MNI) |  |  | Cluster size<br>(voxels) | Z-score<br>at peak | Corrected<br>P-value |
| --- | --- | --- | --- | --- | --- | --- |
|  | X | Y | Z |  |  |  |
| **<u>Feedback On</u>** |  |  |  |  |  |  |
| *L Putamen* | -15 | 7 | -7 | 2252 | 6.94 | P << 0.01 |
| *L Angular Gyrus* | -53 | -71 | 49 | 357 | 4.95 | P < 0.01 |
| *Mid Orbital Gyrus* | -9 | 51 | -3 | 290 | 5.10 | P < 0.01 |
| **<u>Feedback Off</u>** |  |  |  |  |  |  |
| *R Putamen* | 25 | 9 | -5 | 1441 | 5.56 | P << 0.01 |
| *L Putamen* | -23 | 7 | 9 | 1061 | 5.98 | P << 0.01 |
| *R Inferior Parietal Lobule* | 57 | -43 | 55 | 1027 | 4.58 | P << 0.01 |
| *L Inferior Parietal Lobule* | -55 | -49 | 51 | 500 | 4.61 | P << 0.01 |
| *R Middle Frontal Gyrus* | 37 | 35 | 21 | 342 | 5.51 | P < 0.01 |
| *L Cerebellum VIII* | -31 | -67 | -51 | 284 | 5.10 | P < 0.01 |
| *L Cerebellum (Crus 2)* | -13 | -81 | -31 | 256 | 5.44 | P < 0.01 |
| *R Inferior Frontal Gyrus* | 49 | 17 | 9 | 245 | 4.68 | P < 0.01 |
| *R Superior Medial Gyrus* | 7 | 27 | 45 | 233 | 4.57 | P < 0.01 |
| *R Precentral Gyrus* | 45 | -3 | 39 | 206 | 4.19 | P < 0.01 |
| *R Superior Frontal Gyrus* | 25 | 9 | 55 | 201 | 4.67 | P < 0.01 |
| *R Inferior Frontal Gyrus* | 41 | 45 | -1 | 187 | 4.86 | P < 0.01 |
| **<u>Feedback On (Test)</u>** |  |  |  |  |  |  |
| *R Putamen* | 29 | 5 | 1 | 185 | 4.77 | P < 0.01 |
| *L Putamen* | -29 | 9 | -5 | 184 | 4.96 | P < 0.01 |
| *\*Mid Orbital Gyrus* | -1 | 51 | -3 | 28 | 3.79 | N.S. |
| **<u>Random Motor Control</u>** |  |  |  |  |  |  |
| *L Sensorimotor Cortex (S1/M1)* | -51 | -21 | 47 | 4671 | 7.00 | P << 0.01 |
| *R Cerebellum (VI)* | 17 | -53 | -21 | 2588 | 7.25 | P << 0.01 |
| *R Inferior Parietal Lobule* | 41 | -45 | 55 | 661 | 4.86 | P << 0.01 |
| *L Cerebellum (VI)* | -33 | -63 | -25 | 512 | 5.20 | P << 0.01 |
| *L Thalamus* | -17 | -23 | 9 | 306 | 6.34 | P < 0.01 |
| *L Putamen* | -31 | -5 | -1 | 186 | 5.44 | P < 0.01 |

\*voxel-wise $p < 0.005$ for this cluster presented in Figure 2A, $p < 0.001$ was used for all the others clusters.

### ROI analysis

To analyze activity in the putamen and caudate nucleus, we defined ROIs using the intersection of the Harvard-Oxford subcortical structural atlas used in FSL (https://fsl.fmrib.ox.ac.uk/fsl/fslwiki/Atlases) and the Talairach Daemon database used in AFNI. For the caudate nucleus, it was further subdivided into the “head” and “body” regions based on the AFNI’s atlas. For the vmPFC ROI, we used the Talairach Daemon database. As suggested by the literature ^15^, the putamen ROIs were further divided into the anterior (Y > 0) and posterior (Y < 0) regions with a 1-voxel gap between them to reduce partial volume effects. We resampled both atlases to 2-mm-cubic voxels matched to the spatial resolution of acquired functional images. Finally, for ROC analysis, we defined the whole striatum combining the caudate nucleus and putamen from both atlases.

For each ROI, The *NiftiLabelsMasker* function of the *Nilearn* Python module was used to extract the average *β* estimates from the parametric GLM analysis of the success rate for each ROI. The extracted data were analyzed using a two-way repeated measures ANOVA with the region (anterior and posterior, left and right) for each scan session (main and localizer) as within-subject factors. Subsequently, its effect sizes were estimated utilizing partial *η*-squared values, and Greenhouse-Geisser correct were performed.

### Striatum topography of task fMRI vs. anatomy

We assessed the spatial agreement of the anatomically defined striatum and the region identified as significant by the group-level GLM analysis using a parametric regressor. The voxel-wise *β* values for each participant were first converted to z-scores using *3dttest++* with the “-toz” option, after which the voxel-wise threshold of the z-scores was applied to create an ROC curve. The sensitivity and specificity indicate the proportion of voxels within the anatomically defined striatum (i.e., true positive) and outside the striatum smaller (i.e., true negative) than a varying threshold of the z-score. The area under the ROC curve (AUC) ranging from 0 to 1 indicates the performance of a classifier. The AUC of a perfect classifier is equal to 1, and that of a random classifier is equal to 0.5. Although these procedures varied slightly between individuals compared to anatomical volumes, the domain’s boundaries were clearly delineated at the group level.

## Results

### Behavioral data analysis

Twenty-four participants completed the experiment, which lasted about an hour. As described in Materials and Methods, they freely moved and stopped their right fingers following a message that appeared on a computer screen, “Move” and “Stop” (Figure 1A). In the main session, they learned to reach an on-screen target that appeared in one of four corners with a cursor controlled by right fingers, of which movement was recorded by a data glove (Figure 1B). Participants performed the task with or without continuous online visual feedback of the cursor position while a target turned red when reached with a cursor in both feedback conditions. They gradually learned to reach a target across the alternating conditions (Figure 1C).

The initial and final performances roughly matched those of our previous study with a similar experiment^14^. As expected, learning performance was much higher when the cursor movement was visible, suggesting that participants heavily relied on online visual feedback to perform the complicated motor task. A two-way repeated measures ANOVA found a significant main effect of the learning stage (Early, block 1-24, i.e. total 24 blocks; Late, block 25-48, i.e. total 24 blocks; *F*(1,23) = 70.98, *p* < 10^−4^) and feedback conditions (cursor on vs. off, *F*(1,23) = 227.1, *p* < 10^−4^), and their interaction *F*(1,23) = 101.8, *p* < 10^−4^). Although the effect of the learning stage was much less in the discrete condition, participants improved their performance without online cursor feedback (Early, block 2, 4, 6, ⋯, 24, i.e. total 12 blocks; Late, 26, 28, 30, ⋯, 48, i.e. total 12 blocks; *T*(23) = 4.37, *p* < 0.001), substantiating the contribution of proprioceptive feedback to learning. When the set of target locations suddenly altered to the other one (Figure 1B) in the test blocks with available online cursor feedback, the performance did not change (block 37, 39, 41, ⋯, 47, i.e. total 6 blocks; block 49-60, i.e. total 12 blocks; in Figure 1C, *T*(23) = 0.77, *p* = 0.45). This result suggests that the participants learned to map between finger and cursor movements rather than simply memorizing the hand postures corresponding to targets^14^.

### Corticostriatal responses modulated by learning performance

We hypothesized that the red color indicating “success”, to which a target changes when virtually reached in both feedback conditions, would intrinsically motivate participants to improve their performance without external monetary reward^14^. Thus, to identify brain regions related to this goal-directed motor control, we first computed the task performance as a proportion of time turning on red for each trial, every 4 seconds. Then, it was convolved with a canonical hemodynamic response function to construct a parametric regressor used in general linear model (GLM) analysis, which identified regions where performance modulates activities (Table 1, Figure S1). Due to the task instruction that participants should stop moving their fingers once a target is reached, the parametric regressor was negatively correlated with the extent of movement. Thus, we found various regions related to motor control, such as sensorimotor cortices, thalamus, parietal regions, and cerebellum. Here, we focused on the regions where performance positively modulated activities (Table 1).

We separately performed this analysis for each of the two feedback conditions and a test condition and then compared the results (Figure 2). For both feedback conditions, we primarily found robust positive modulation in the bilateral striatum and early visual cortex (Figure 2 and Table 1). The striatum is the only region where we identified clusters larger than 450 voxels at the highly stringent threshold of *p* < 5.0 × 10^−5^. This result motivated us to analyze further the sensitivity of the GLM in delineating the anatomically defined striatum. For stepwise increasing z-thresholds [z(min)=-8, z(max)=8, z(step)=0.5], an ROC analysis found that the voxels in the striatum are highly selective for our GLM method compared to the rest of the brain regions, resulting in AUC (Area Under Curve) scores of 0.95 (cursor-on), 0.92 (cursor-off) (Figure 2B). The more robust response in the striatum than in the visual cortices further supports the hypothesis that the visual feedback of the performance would convey intrinsic reward to participants. To demonstrate how brain activity relates to performance, we illustrated the correlation between fMRI signals in the striatal region and a parametric regressor that modulates performance. (Figure S2).

**Figure 2.**
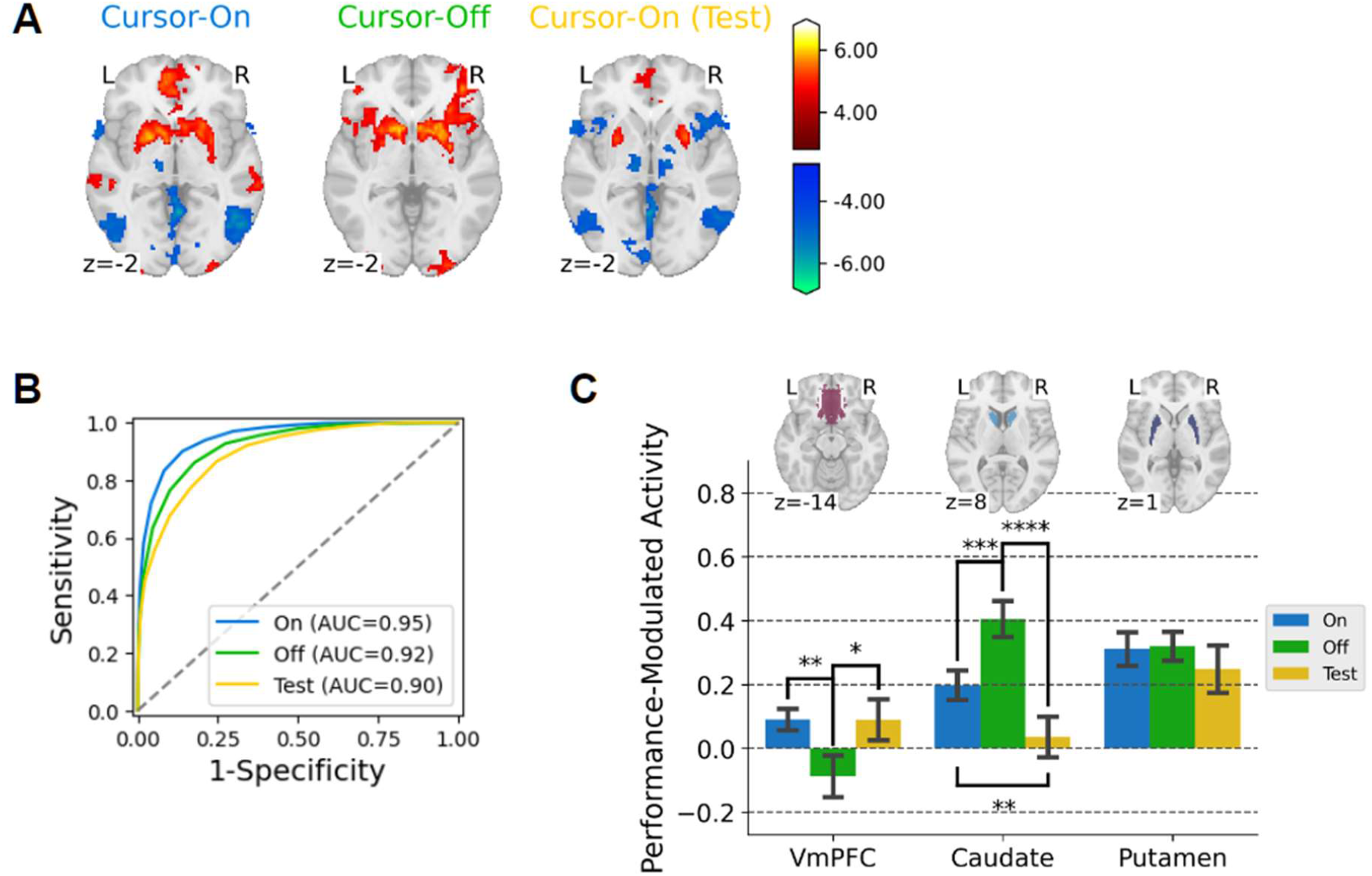
Brain regions where activity is modulated by performance in three learning conditions, “cursor-on” (blue), “cursor-off” (green), and “cursor-on (test)” conditions. (A) The results of whole-brain voxel-wise GLM analysis with parametric modulation were converted to z-scores, and clusters revealed significant positive modulation (voxel-wise p<0.005 for a visualization purpose). (B) The receiver operating characteristic (ROC) curves for the three conditions show highly selective activity within an anatomically defined striatum compared to the whole brain. (C) Comparison of activity in the corticostriatal ROIs (vmPFC, caudate nucleus, and putamen) among the three conditions.

**Figure 3.**
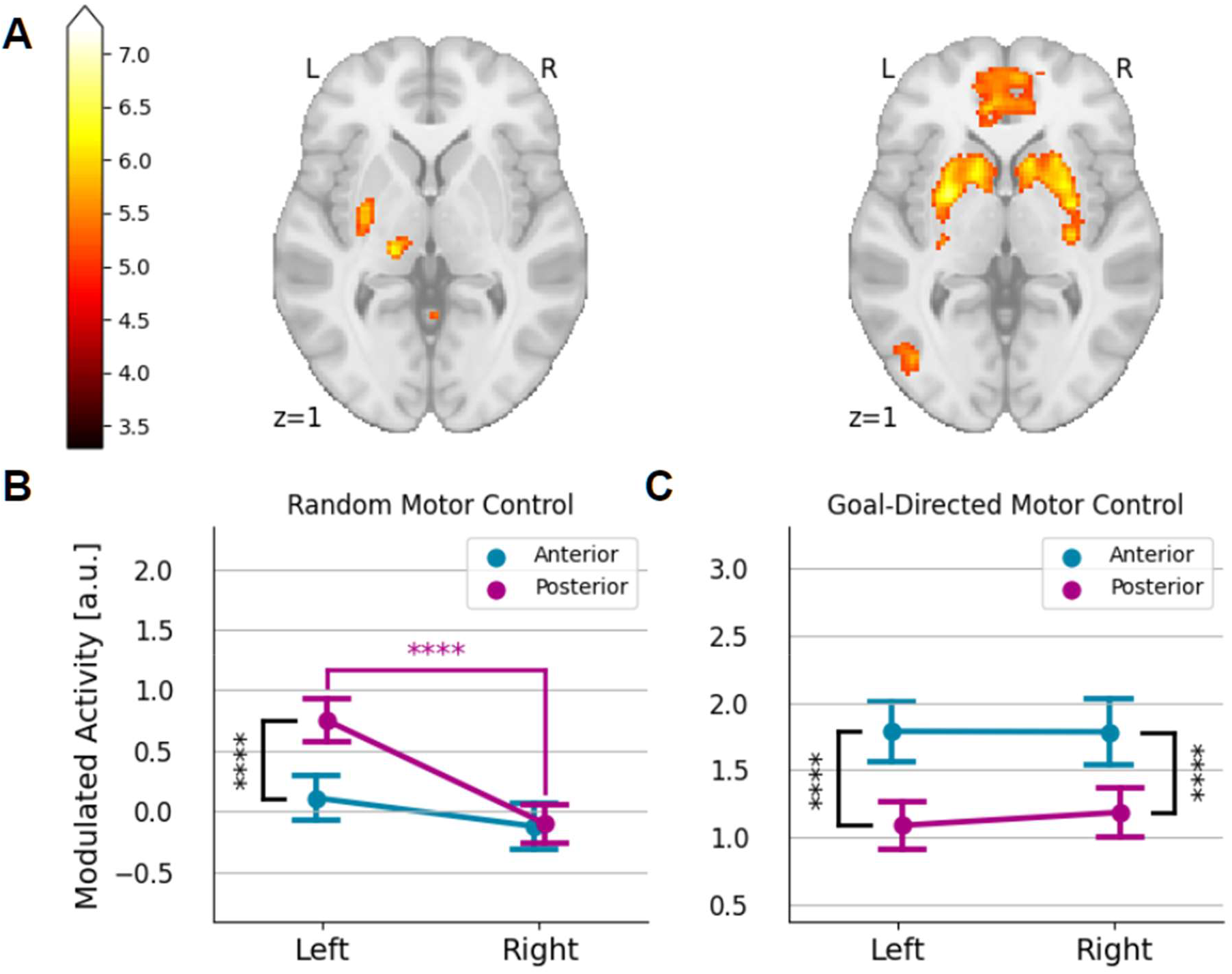
(A) Comparison of voxel-wise striatal responses related to performance during goal-directed motor control (right) and those related to random motor control (left). (B) During random motor control, the left posterior putamen showed contralateral activation. (C) Significant activations of the bilateral anterior putamen were observed for the performance-modulated response during goal-directed motor control. Error bars indicate SEM. *p<0.05; **p<0.01; ***p<0.001; ****p<0.0001 (uncorrected p).

While the overall striatal responses are similar for both feedback conditions, there were prominent differences between the conditions. First, activity in the middle temporal region (V5) is negatively modulated by the performance only when online cursor feedback is available. Since the performance is lower with larger cursor movement, the performance-modulating regressor found the V5 region negative, which is sensitive to the visual motion of the cursor. Thus, this activity was not found for the condition without online cursor feedback (cursor-off condition). In contrast, the activity in the anterior insular cortex, an important region of the salience network^16^, is found only for the “cursor-off” condition. In the “cursor-off” condition, hitting a target was more difficult and thus the visual feedback provided as a “red signal” garnered increased attention, leading to increased activity within the salience network. Likely for the same reason, the insular cortex’s activity was negatively influenced during the test condition with reduced saliency, where the online cursor remained consistently accessible without interleaved “cursor-off” blocks.

Interestingly, there was a clear difference between the two feedback conditions in the ventromedial prefrontal cortex (vmPFC) (Figure 2A). An ROI analysis revealed that the activity was significantly higher in the “cursor-on” condition than in the “cursor-off” condition (*T*(23) = 3.04, *p* < 0.01), without significant activity for the “cursor-off” condition (*T*(23) = 1.33, *p* = 0.20) (Figure 2A, C). In contrast, in the “cursor-off” condition, the caudate nucleus activity was more robust than in the “cursor-on” condition (*T*(23) = 3.80, *p* < 0.001) (Figure 2C). We also found a significant interaction of the feedback condition and subregions (caudate and putamen) (*F*(1,23) = 26.0, *p* < 10^−4^, 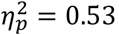). The caudate and putamen regions respectively exhibited more pronounced activity in the “cursor-off” (*T*(23) = 2.52, *p* = 0.019) and in the “cursor-on” conditions (*T*(23) = 3.71, *p* = 0.0015). However, there was no significant difference in the putamen between the feedback conditions (*T*(23) = 0.16, *p* = 0.87). These dissociable corticostriatal responses would implicate distinct roles of the vmPFC and the striatum in learning action values from feedback^17-21^.

The corticostriatal responses for the test condition in which the cursor feedback was similar to the “cursor-on” condition with highly selective response in the striatum with an AUC score of 0.90 (Figure 2B) and negative modulation in V5 region due to online cursor feedback (Figure 2A). This was expected since the online cursor feedback was also available in the test condition. However, an ROI analysis revealed that the overall level of the striatal activity decreased compared to the other two feedback conditions only in the caudate nucleus (vs. cursor-on: *T*(23) = 2.88, *p* < 0.01; vs. cursor-off: *T*(23) = 5.22, *p* < 10^−4^) contributing to lowered AUC score (Figure 2B, C). The activity in the vmPFC for the test condition was comparable to the “cursor-on” condition (*T*(23) = 0.017, *p* > 0.5) and higher than the “cursor-off” condition (*T*(23) = 2.12, *p* < 0.05) (Figure 2C), although the cluster in the vmPFC did not survive after cluster-level correction (Table 1). The decreased activity in the caudate nucleus is potentially due to the learning effect, as shown in our previous study^14^. Although the target locations were altered in the test condition, participants learned the identical mapping throughout the session. The comparable activity in the vmPFC to the “cursor-on” condition is also consistent with the previous study (Figure 2C). However, the extent of the vmPFC response is relatively limited for the test condition (Table 1).

### Corticostriatal responses modulated by random finger movement

As we noted previously, the performance of our task is negatively related to the amount of finger movement due to the goal of the task staying longer in the target. Thus, instead of controlling motor control components in a GLM analysis, we analyzed the dataset from a separate experiment where participants randomly moved their fingers without any task-related visual feedback. Through this experiment, we aimed to confirm that the response patterns associated with performance (Figure 2) were unique compared to those connected to motor control. For this separate dataset, we employed a standard GLM analysis to contrast the “Move” and “Stop” conditions (see Materials and Methods for more details). This simple contrast roughly aligned with the GLM analysis using a parametric regressor for the amount of finger movement since participants moved their fingers constantly during “Move” and stopped during “Stop”. The analysis identified well-known regions related to motor function. These include motor and somatosensory cortices (M1/S1), supplementary motor area (SMA), left thalamus, inferior parietal cortex, insula, left posterior putamen, and cerebellum lobules 6 and 8 (Table 1). We also confirmed the typical laterality of the motor system, that is, contralaterally dominant in M1/S1, thalamus, and putamen and ipsilaterally dominant in the cerebellum.

In contrast to the performance-modulated response during the goal-directed finger movement discussed earlier, the striatal activity during random finger movement was less pronounced, particularly due to the lack of significantly positive activity in the caudate nucleus although they are significantly negative (left: *T*(23) = 2.71, *p* = 0.012, right: *T*(23) = 2.34, *p* = 0.028) (Figure 3A, C). Thus, the overall response to random finger movement was also higher in the putamen than in the caudate nucleus (left: *T*(23) = 6.49, *p* < 10^−5^, right: *T*(23) = 3.07, *p* < 0.01). A subsequent ROI analysis for putamen subregions (Figure 3B) discovered significant interaction of anteriority and laterality with distinct activity in the left posterior putamen only (anteriority, *F*(1,23) = 18.70, *p* < 0.001,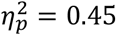; laterality, *F*(1,23) = 35.29, *p* < 1 × 10^−5^,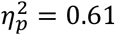; interaction, *F*(1,23) = 19.28, *p* < 0.001, 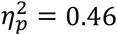). The activity in the posterior putamen is contrasted with the performance-modulated activity, which is more robust in the anterior putamen (*F*(1,23) = 36.19, *p <* 10^−5^, 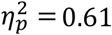) (Figure 3C) regardless of laterality and feedback condition (combined “cursor-on” and “cursor-off” in Figure 3C and separated in Figure S3), which is also the case for the anterior versus posterior caudate nucleus (*F*(1,23) = 11.65, *p <* 0.01,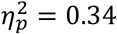). The result is consistent with previous studies reporting the role of contralateral posterior putamen in a simple motor execution^22-24^ and the positive performance-related activity in the anterior striatum during the early stage of learning^14^. Similar to the putamen activity, the caudate activity that modulates performance was also found to be more robust in the anterior region than in the posterior region (*T*(23) = 3.27, *p* < 0.01) regardless of laterality (We provided a full result of the striatal subregion ROI analysis in Figure S3.

## Discussion

We designed a visuomotor learning task with two interleaved visual feedbacks, continuous (online cursor) and discrete (offline cursor), to understand the role of visual feedback in motor skill learning. Our GLM analysis with a parametric regressor modulating participants’ performance provided by both visual feedbacks revealed robust activity in the striatum with strikingly high sensitivity and specificity. The entire region of the anatomically defined striatum was highly responsive to visual feedback.

The effects of visual feedback on the striatal response were spatially distinct from those of motor control, which were mapped on the anterior striatum and contralateral posterior putamen, respectively. This conclusion is corroborated by well-established parallel and segregated cortico-striatal connections comprising the anterior cognitive loop and posterior sensorimotor loop. Random finger movements without visual feedback did not elicit the global response in the striatum, but rather the local response in the contralateral (left) posterior putamen ^25,26^, which is predominantly interconnected with the primary motor cortex ^27,28^. This result is also consistent with previous studies reporting a gradual shift of striatal activation from the anterior to the posterior region ^9,14^. In other words, the anterior region is initially involved in the goal-directed movement with reward feedback, whereas the posterior region is associated with more habitual movement independent of reward feedback in the late stage of learning. Due to limited practice in a scanner lasting less than an hour, the response to the visual feedback was higher in the anterior striatum. In contrast, the response to random finger movement, which is not goal-directed but rather habitual in the absence of feedback, was more significant in the posterior striatum, specifically in the contralateral putamen. This observation, in conjunction with our currently obtained results from different cortical motor areas, supports the putamen’s predominant role as a motor hub within the striatum, highlighting its contrasting relationship with the caudate nucleus. ^29-31^.

We found a double dissociation of corticostriatal activity between two feedback conditions. In the presence of online cursor feedback, the vmPFC exhibited more significant activity related to performance, while the caudate nucleus demonstrated reduced activity compared to the condition lacking online feedback. We speculate that the distinction is closely related to model-based versus model-free reinforcement learning^32-34^. Particularly, we conjecture that error-based learning occurs when the cursor is visible, whereas reward-based learning takes place when the cursor is not visible. Specifically, a forward model, which is a finger-to-cursor mapping, is learned only in the “cursor-on” condition while it is retrieved in the “cursor-off” condition. In the reinforcement learning framework, the forward model provides how an agent’s state (i.e., hand posture and cursor position) is transitioned, a state-transition rule. Our results clearly indicate that participants did not merely memorize hand postures for targets; instead, they learned the mapping between finger and cursor movements.

Thus, the vmPFC activity in the “cursor-on” condition would be related to the state-action value predicted from the forward model in model-based reinforcement learning as supported by previous studies using decision-making tasks^17-21,35^. However, when the online cursor feedback is unavailable, participants should rely more on trial-and-error than an uncertain forward model without online feedback and thus simply reinforce actions associated with larger rewards in a model-free manner. The activity in the striatum would be contributed to model-free learning^32,34^.

Regarding the caudate nucleus activity, the lower prediction of success in the “cursor-off” condition and thus larger reward prediction errors when hitting a target would be related to the higher activity compared to the “cursor-on” condition. Indeed, previous research has shown that the caudate nucleus has a larger role in predicting reward errors, similar to the ventral striatum’s function. Conversely, the putamen is predominantly engaged in predicting rewards through the learning of stimulus-action-reward associations in a more certain condition, leading to a more habitual behavior^36,37^. Thus, in the “cursor-off” condition with larger uncertainty, the caudate nucleus would be more sensitive to reward prediction errors. This interpretation aligns with the decreased caudate activity in the test condition, a result of learning with reduced uncertainty. To test this idea, it is necessary to develop computational models that can predict action values and associated reward prediction errors. However, creating such models for the complex motor tasks presented in this study poses a significant challenge^38^.

The highly sensitive striatal response appears to be related to intrinsic reward, motivating to learn the complicated motor skill learning in the absence of a monetary incentive. The intrinsic reward for good performance is sufficient to elicit striatal activity^39-41^, while specific subregions of the striatum are dissociable depending on the nature of the reward, extrinsic versus intrinsic^41^. The extrinsic and intrinsic rewards have dissociable effects on motor learning. The former influences early rapid improvements in speed and accuracy, whereas the latter influences training-based enhancement^42^. However, the extrinsic monetary reward could undermine the intrinsic reward processing, lowering motivation or performance^43,44^. It would be fascinating to determine if the extrinsic reward inhibits or enhances the effect of intrinsic reward on motor learning, including long-term retention of motor memory.

There are several limitations in the current study, primarily due to the experiment design and the motor task. First, our main GLM analysis using a parametric regressor did not remove the effects of motor components, such as kinematics of finger movements on the corticostriatal activity, since participants were instructed to stop moving fingers when a target is reached with a red signal. Due to this instruction, the extent of finger movement is negatively correlated with performance, which we used for a parametric regressor. Consequently, it was difficult to remove the confounding motor control effects in the main analysis. Moreover, it is also hard to reject an alternative hypothesis that the red signal might play a role as an instruction instead of performance feedback. Thus, future studies should test another condition with online cursor feedback but without the red signal to fully understand the respective role of the two components of the visual feedback, the cursor and the red signal. Despite these limitations, our main results of corticostriatal response patterns are more likely related to goal-directed learning based on visual feedback because we found distinct corticostriatal activities in a separate simple motor control task. This result is consistent with our previous study using a similar experiment design^14^.

Lastly, our findings suggest a crucial role of immediate performance feedback in eliciting striatal responses. If this association extends to dopamine release, it could potentially aid in restoring the compromised striatal dopamine system in PD patients. Furthermore, it would be even more advantageous for rehabilitation to utilize extrinsic reward or augmented feedback to the extent that it does not undermine the effect of performance feedback^45^. Previous fMRI studies in other cognitive domains have shown that extrinsic^43^ and intrinsic reward^46^ strengthen long-term memory via dopamine release. Together, designing visual feedback directly related to performance would be essential to improve the long-term retention of motor memory, maximizing the treatment effects for PD patients.

## Code availability

All the codes necessary to generate results and figures shown in this study are available upon request.

## Acknowledgments

Neuroimaging was performed at the Center for Neuroscience Imaging Research located in Sungkyunkwan University, supported by Institute for Basic Science.

