## Supplemental Table 1, Supplemental Figure S1, Supplemental Figure S2, Supplemental Figure S3 for "Effects of visual feedback on corticostriatal activity during continuous *de novo* motor skill learning"

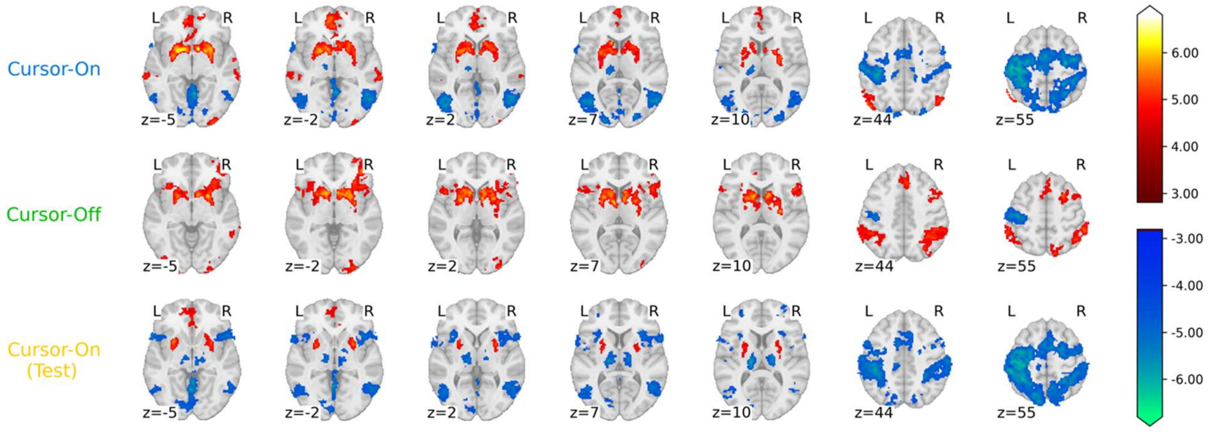


Figure S1. Whole-brain voxelwise GLM analysis for three learning conditions with "cursor-on", "cursor-off", and "cursor-on (test)".

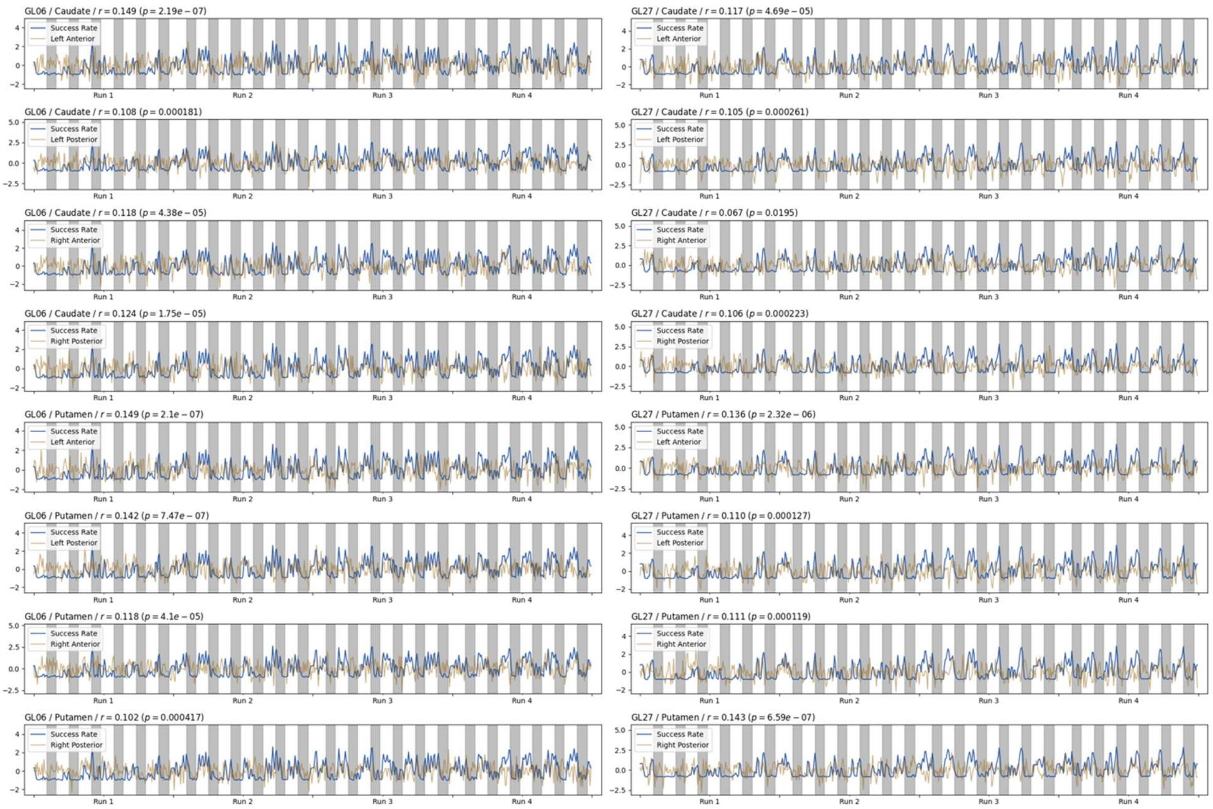


Figure S2. Examples of a parametric regressor, which is a HRF-convolved trial-by-trial success rate, (blue) and fMRI BOLD signal (gold) extracted from a voxel with a peak response in the caudate nucleus and putamen for two representative participants. Six regressors of non-interest related to the head motion and 4th order polynomial trends were projected out from the BOLD signals. Then the BOLD signals were resampled every 4 s (original TR = 2 s) for better comparison with the parametric regressor that modulates trial-by-trial success rate.

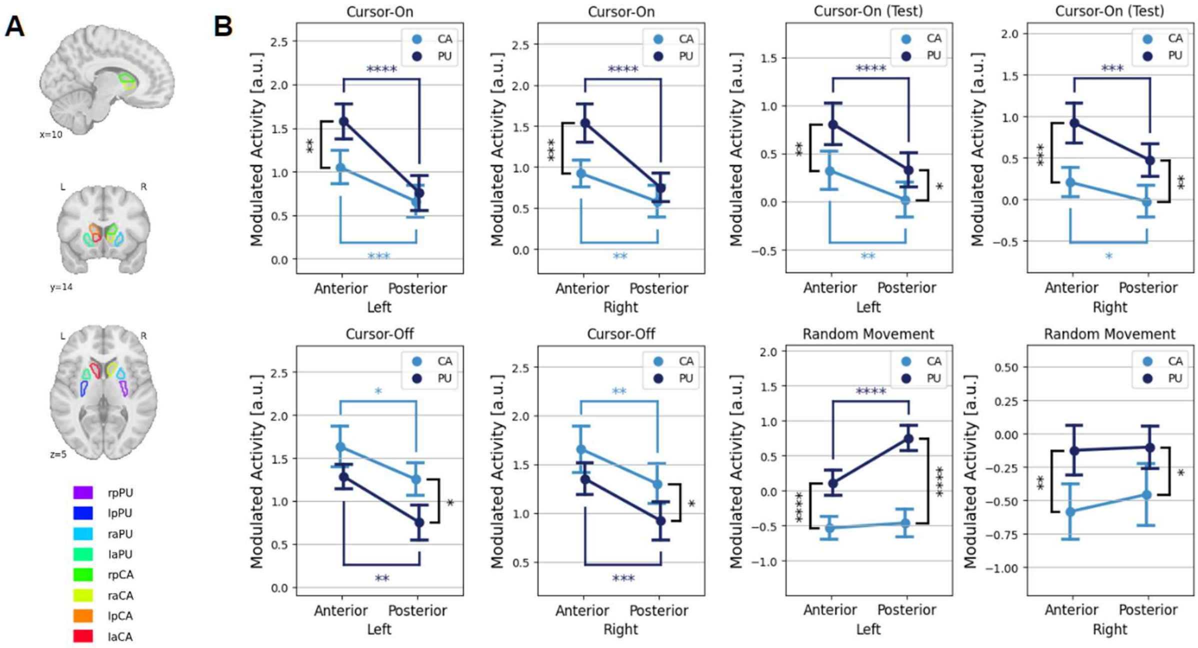


Figure S3. (A) As sub-regions of the striatum, the putamen and caudate were independently split into anterior and posterior portions. Where r stands for right, l for left, a for anterior, and p for posterior. (B) Spatial dissociation of the striatal activity for four different conditions, three conditions for performance-modulated activity "Cursor-On", "Cursor-Off", "Cursor-On (Test)", and one condition for random motor control. Error bars indicate SEM. *p<0.05; **p<0.01; ***p<0.001; ****p<0.0001 (uncorrected p).
